# Emergence of CC17 vancomycin variable *Enterococcus faecium* in India

**DOI:** 10.1101/2022.07.29.501338

**Authors:** Madhan Sugumar, Sreeram Chandra Murthy Peela, Lakshmi Shree Viswanath, Kamini Walia, Sujatha Sistla

## Abstract

Vancomycin variable enterococci (VVE) are those isolates that are susceptible to glycopeptide like vancomycin but harbour the resistance determinants like vanA gene. Globally the emergence of such strains was reported, and from India our centre was the first to report them. The present study focussed on analysing genome content of these strains from India to assess their genetic diversity. While the five isolates belonged to three sequence types, all the isolates were of clonal complex CC17. This supports that VVE from India are genetically closely related and are emerging from a single clone.

## Introduction

Enterococci are predominantly non-pathogenic gastrointestinal commensal bacteria that occasionally cause human infections such as septicemia, urinary tract infections, endocarditis, and infection in patients with indwelling catheters, predominantly as opportunistic infections (1,2). Among them, *Enterococcus faecalis* and *Enterococcus faecium* represent the species that account for most infections. *E. faecium* has been able to adapt to the hospital environment, emerging during the last few decades as a leading cause of health-care infections worldwide, and becoming the most challenging species to treat.

The emergence of vancomycin-resistant enterococci (VRE) has been associated with an increase in multidrug-resistant nosocomial infections. Genome plasticity, the presence of multiple antibiotic resistance determinants, and the lack of therapeutic options have contributed to the adaptation of *E. faecium* to hospital environments (3,4). Moreover, high recombination rates and the acquisition of mobile elements in the genome of *E. faecium* have also driven this evolutionary process (5). In terms of antibiotic resistance, one of the most relevant traits acquired by enterococci is resistance to vancomycin due to the *van* gene clusters (6). Furthermore, vancomycin-resistant *E. faecium (VREfm)* frequently exhibits resistance to ampicillin and high-level resistance to aminoglycosides.

The glycopeptide resistance in *E.faecium* is known to be mediated predominantly via *vanA* gene. This gene mediates high level resistance by altering the binding site, and acts along with *van*H, *van*S, and *van*X genes (7). It is interesting to note that in the recent years, *E.faecium* isolates that were phenotypically susceptible to vancomycin, but harbouring the vanA gene were isolated in a few parts of the world.

Our centre was the first to identify and report VVE from India (8).Among the 340 isolates collected across the country, 5 isolates harboured the *vanA* gene in which one isolate was harboured *vanRS* genes. The goal of the current study was to identify the genetic relatedness of these five VVE isolates.

## Methods

### Bacterial isolates

Our hospital is one of the nodal centres involved in characterising *E. faecium* isolates across the county as part of the Antimicrobial Resistance Surveillance Network program (AMRSN) under the Indian Council of Medical Research. As part of the National Antimicrobial Resistance Surveillance Network (AMRSN), five VVE isolates were identified from 340 *E. faecium* isolates from 20 centres across India. These five isolates were selected for genome sequencing

### DNA purification and WGS

The DNA from the five isolates was extracted and purified using QiaAmp DNA Blood Mini Kit following manufacturer instructions for purification of DNA from Gram positive bacteria. About 200ng of purified DNA was used for library preparation using TruSeq Nano DNA library preparation kit. These libraries were loaded onto the Novaseq 6000 platform in paired end mode and with a length of 151bp.

### Genome analysis

The raw FASTQ files were assembled using GHRU SPAdes assembly workflow with inclusion of parameter --careful for complete genome sequences using default values (9). The generated contigs were annotated using Prokka (10). The presence of various genes in the van locus was tested in each genome using LS-BSR method and a score of > 0.9 indicated the presence of the corresponding gene (11). The reference *van* genes were downloaded from NCBI AMR database. The multi-locus sequence typing (MLST) was performed using the MLST tool (12,13).

## Results and Discussion

The five genomes had a mean length of 2.9Mb consistent with the expected genome size of *E.faecium.* When the genome sequence results from 5 isolates were analysed for the presence of genes in the *van* locus, *vanA* genes were detected only in 2 isolates (227150 and 272181) using LS-BSR score cut-offs. The three isolates in which *van*A was not detected by WGS were re-confirmed as VVE using PCR and Sanger di-deoxy sequencing of the amplified products. Genetic relationships by whole-genome and MLST scheme predicted the most common clonal type were (ST80 and ST18) (Table 1). Two of the three isolates from Mumbai were of same ST (ST80). Only one isolate produced *vanR* and *vanS* genes and was identified as ST31. While all these five isolates belonged to three different STs, they belonged to the same clonal complex CC17. This clonal complex is known to be associated with nosocomial infections with increased risk of morbidity and mortality (14). Recent evidence showed that ST80 and ST18 were predominantly VRE (22/51, 43.1%) and can be isolated from stool specimens (15). In another study from Denmark, ST80 accounted for 22% of VVE/VRE in 2015, but was replaced by ST1421 in 2019 (16). This highlights the importance of continued surveillance of VVE in India. Further large-scale screening for VVE from India is underway and is a major thrust of the antimicrobial resistance surveillance network (ICMR-AMRSN).

**Table 1:** MLST types for the five VVE isolates. CC-Clonal Complex, ST – Sequence Type

| Sample ID | Specimens | CC | ST | atpA | ddl | gdh | purK | gyd | pstS | adk |
| --- | --- | --- | --- | --- | --- | --- | --- | --- | --- | --- |
| 148814 | Blood | 17 | 80 | 9 | 1 | 1 | 1 | 12 | 1 | 1 |
| 171861 | Bile | 17 | 80 | 9 | 1 | 1 | 1 | 12 | 1 | 1 |
| 148816 | Blood | 17 | 18 | 7 | 1 | 1 | 1 | 5 | 1 | 1 |
| 227150 | Blood | 17 | 18 | 7 | 1 | 1 | 1 | 5 | 1 | 1 |
| 272181 | Ear swab | 17 | 31 | 1 | 3 | 1 | 1 | 1 | 1 | 3 |

## Conclusions

The vancomycin variable enterococci (VVE) from India are clonally related and belonged to the clonal complex CC17. Further large-scale screening of VVE isolates from India and identifying clonal relationships of VVE may help in understanding the spread of this clonal type in India. Clinical problems such vancomycin resistance detection, surveillance, and horizontal transmission are made worse by the features of the VVE isolates with altered resistance phenotypes. In light of this finding it may be advisable to use both phenotypic and genotypic tests for detection of vancomycin resistance in enterococci.

